# A gel-top model for characterizing mesenchymal features of glioblastoma cells

**DOI:** 10.1101/2025.04.03.647072

**Authors:** Yuanzhong Pan, Priscilla Chan, Jeremy N. Rich, Steve A. Kay, JinSeok Park

## Abstract

Glioblastoma (GBM) remains one of the most aggressive adult cancers, with limited treatment options due to an incomplete understanding of its biology. GBM exhibits different transcriptional subtypes, among which the mesenchymal (mGBM) variant is associated with a particularly aggressive nature and poor outcomes. Developing models that capture mGBM phenotypes could offer new insights into its biology and therapeutic vulnerabilities. However, the lack of in vitro models for mGBM has posed challenges in investigating the aggressive GBM subtype.

Here, we present a *gel-top culture* model in which GBM cells exhibit MES features, including distinct transcriptomic profiles, invasive phenotypes, cell morphology, and cytoskeletal organization. Furthermore, co-culturing GBM cells with microglia in this system revealed enhanced microglial recruitment and interaction, effectively recapitulating the tumor microenvironment of mGBM that traditional 2D culture fails to model.

Using the bioinformatic tool, Small Molecule Suite, we identified casein kinase 2 (CK2) as a targetable molecule in GBM cells within our system, potentially driving MES features. Notably, CK2 inhibitors reduced the mesenchymal signature of mGBM cells, suggesting their potential as a targeted therapy for mGBM.

Together, our findings suggest that the gel-top culture model replicates key features of mGBM, serving as a valuable platform for studying its biology and identifying novel therapeutic strategies.

## Introduction

Glioblastoma (GBM) is the most aggressive and common form of malignant primary brain tumor in adults^1,2^. Despite advances in treatment, including surgery, radiation, and temozolomide-based chemotherapy, GBM remains incurable, with a five-year survival rate of less than 10%^2,3^. A major challenge in treating GBM is the limited understanding of its complex biology, including its highly heterogeneous cellular composition and adaptive resistance mechanisms^4^.

Molecular profiling has identified distinct subtypes reflecting the heterogeneity of GBM. The analysis of the TCGA cohort and subsequent studies defined three subtypes: proneural, classical, and mesenchymal^5^. These subtypes have significantly improved our understanding of the GBM molecular landscape and emphasized its intrinsic plasticity, allowing transition along the spectrum of different subtypes. This insight has led to the development of therapeutics aimed at repressing the transition toward inferior subtypes^6^. Such efforts require a better understanding of the different GBM subtypes and their response to treatments. Among all the subtypes, mesenchymal GBM (mGBM) consistently exhibits enhanced invasiveness, greater resistance to therapy, and poorer prognosis^7,8^. To develop potential therapeutics for this aggressive type, it would be essential to understand the underlying biology of mGBM.

One of the key challenges in investigating mGBM biology is the lack of an appropriate phenotypic model that effectively recapitulates its distinct cellular features. Traditional 2D models are physiologically limited and have failed to capture the spectrum of cell behaviors. Furthermore, current *in vivo* models are relatively expensive, difficult to characterize, and challenging to control for different variables. An ideal model should reflect the key cellular properties of mGBM and provide a robust platform for studying its biology and testing potential therapeutics.

Hydrogels present unique opportunities for developing new models for studying cancer including GBM^9^. Hydrogels can mimic and control different aspects of the natural extracellular matrix (ECM), providing physical support as a scaffold in the tumor microenvironment (TME), and have demonstrated significant potential for studying tumor biology^10^. When cultured in the matrices, cells demonstrate more physically relevant features not observed in traditional 2D culture. However, the phenotypic landscapes of GBM cells cultured in hydrogels and their application, such as drug testing, remain underexplored. Therefore, we leverage a hydrogel-based platform to develop in vitro models that better characterize the properties of GBM cells and facilitate the evaluation of potential therapeutics.

This work tested a gel-top model for characterizing mGBM cells. We demonstrated that GBM cells cultured on the hydrogel model exhibited mesenchymal features, including increased invasiveness and elevated expression of mesenchymal signature genes within a relatively short time frame. Furthermore, this time-efficient model is adaptable for incorporating different components of the TME, such as microglia, effectively replicating the complex TME interactions of mGBM cells and enhancing microglial recruitment. Additionally, using the bioinformatics tool, Small Molecule Suite^11^, we identified casein kinase 2 (CK2) as a potential therapeutic target for repressing mesenchymal features of GBM cells on the hydrogel model. We further confirmed that CK2 inhibitors suppressed the mesenchymal characteristics of GBM cells. Together, our findings suggest that the gel-top culture model replicates key features of mGBM cells, serving as a valuable platform for studying its biology and identifying novel therapeutic strategies.

## Results

### Gel-top culture on BME shows unified and elevated mesenchymal features of GBM cells

GBM cells undergoing mesenchymal transition can often be characterized by their mesenchymal morphology, enhanced spreading and elongation^12^. However, when we culture two mGBM lines on traditional 2D plastic surfaces, they showed different morphology: T387 grew as neurospheres, whereas MGG31 exhibited adherent mesenchymal-like morphology (**Figure 1A**). We re-affirmed the mesenchymal subtype of the two cell lines by scoring for signature genes of different subtypes but found that MGG31 has a higher mesenchymal signature than T387 cells (**Supplementary Figure 1A, B**). This finding suggests that the 2D culture model fails to fully capture the mesenchymal phenotype unless the cells possess a strong mesenchymal identity. To characterize cells along the mesenchymal transition axis, an *in vitro* model that consistently demonstrates unified mesenchymal features is required.

**Figure 1.**
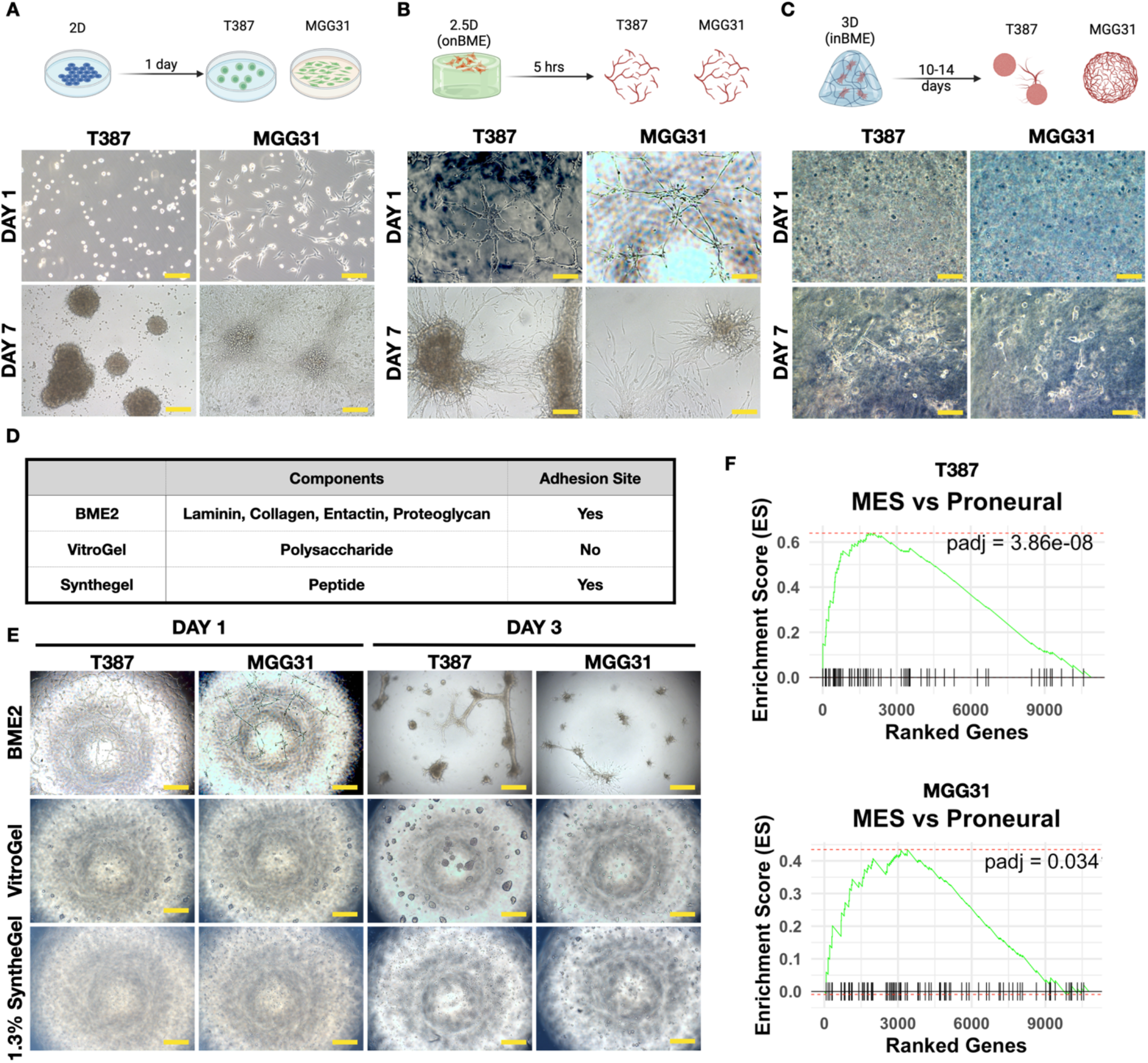
Culture of GBM cells on BME is a time-saving model that exhibits and elevates their mesenchymal signature. **A-C**. Images of two mesenchymal GBM cells under traditional 2D culture (A), onBME culture (B), and 3D inBME culture of on day 1 and day 7. Scale bar = 200 µm. **D**. Summary of the properties of different types of commercially available hydrogels tested for gel-top culture of GBM cells. **E**. Images of cell morphology of GBM cells cultured on top of different materials. Scale bar = 500 µm. **F**. GSEA analysis of highly expressed genes in T387 (top) and MGG31(bottom) on 2.5D (onBME) vs. 2D culture with the gene set mesenchymal versus proneural (see methods).

We tested if applying hydrogel matrices can induce GBM cells to show unified mesenchymal features. Using basement membrane extract (BME), the most commonly used matrix for organoid studies, we generated hydrogel cell culture models: 2.5D (on top of the BME hydrogel, onBME), and 3D (inside the hydrogel, inBME) cultures (**Figure 1B**). Interestingly, when cultured on BME, both GBM cell lines self-assembled into a mesh-like network structure after 6 hours (**Figure 1B**, top panel). The network structure arises from the formation of branched extensions from mGBM cells, indicative of their invasiveness in migrating and interacting with the surrounding ECM. Consistently, after seven days of culture, tumor cells showed extensive invasion into the matrix (**Figure 1B**, bottom panel). On the other hand, 3D cultures showed slow growth and self-assembling (**Figure 1C**). Their self-assembly feature can only be observed after 14 days of culture, and T387 only showed single-cell clones without obvious interaction across different colonies (**Supplementary Figure 1C**). Because 2.5D culture provided a unified phenotype and significantly saved time compared to 3D culture, we chose 2.5D culture for further experiments.

We next examined if the materials of hydrogels play a role in the morphology of GBMs in 2.5D, representing mGBM features. The natural ECM contains various active materials, including proteins (collagen, laminin, etc.) and proteoglycans^13^. Engineered matrices, on the other hand, contain more defined but usually less complex components for different purposes. In addition to BME2, which is a natural composite material, we tested two other commercially available gels that are designer peptide-based (SyntheGel) and saccharide-based (VitroGel, summarized in **Figure 1D**). On the saccharide-based VitroGel, T387 grew as spheres, like when they are cultured without matrices, whereas MGG31 did not show clear proliferation or network formation, indicative of invasiveness (**Figure 1E**, middle panel). There reduced mGBM phenotypes could be because polysaccharide-based gels do not provide adhesion sites, such as the RGD sequence, for the cells to adhere to. Under 2.5D culture, both GBM cells failed to proliferate on the peptide-based SyntheGel (**Figure 1E**, bottom panel). These results suggest that a matrix that contains both peptides and saccharides is necessary for GBM cells to self-assemble and show mesenchymal morphology.

Because onBME culture of the two GBM cells revealed a shared morphological phenotype despite their different initial mesenchymal levels, we hypothesized that onBME culture could elevate the mesenchymal signature of GBM cells. We examined this hypothesis using different scoring metrics. First, we defined a gene set, the differentially expressed genes (DEGs, defined as log2 fold change < -2 and p-value < 0.001, a resulting set of 170 genes) by comparing mesenchymal and proneural subtypes in the TCGA dataset. Gene set enrichment assay (GSEA) on the DEGs comparing onBME and 2D culture showed a significantly enhanced enrichment score (**Figure 1F**). Consistently, GSEA analysis of DEGs comparing onBME and 2D culture on the hallmark EMT gene set, showing significant enrichment in mGBM^14^, showed a significant enrichment score for T387 (Supplementary Figure 1D). The result for MGG31 was not statistically significant, likely because MGG31 is already highly mesenchymal.

We also compared the module score of the GBM cells cultured on 2D versus onBME using the matrix and representation defined by Neftel *et al*^15^. Our results confirmed that MGG31 showed a higher mesenchymal module score than T387, and onBME culture led to a shift of relative module scores towards the mesenchymal state (**Supplementary Figure 1E**). We also performed principal component analysis (PCA) analysis on the transcription data of different cultures (**Supplementary Figure 1F**). Interestingly, the transcriptional differences between 2D and onBME culture are almost only accounted for by the second principal component (PC2), while the first principal component may represent the differences between two GBM cell lines. We therefore ranked the genes by their weights in PC2 and ran GSEA analysis on the mesenchymal signature gene set which showed a significant enrichment score (**Supplementary Figure 1F**). Together, these results provided strong evidence that onBME culture elevates the mesenchymal molecular feature of mGBM cells.

Mesenchymal GBM cells exhibit characteristic features such as high secretion of ECM genes, distinct organization of the cytoskeleton, hypoxic metabolism, and a relatively lower proportion of proliferating cells^8,15^. We reanalyzed the TCGA glioblastoma dataset to find differentially enriched biological processes between the mesenchymal and proneural subtypes. Significantly enriched gene ontology (GO) terms included those related to ECM organization and hypoxia, which may reflect mGBM characteristics (**Figure 2A**)^16,17^.

**Figure 2.**
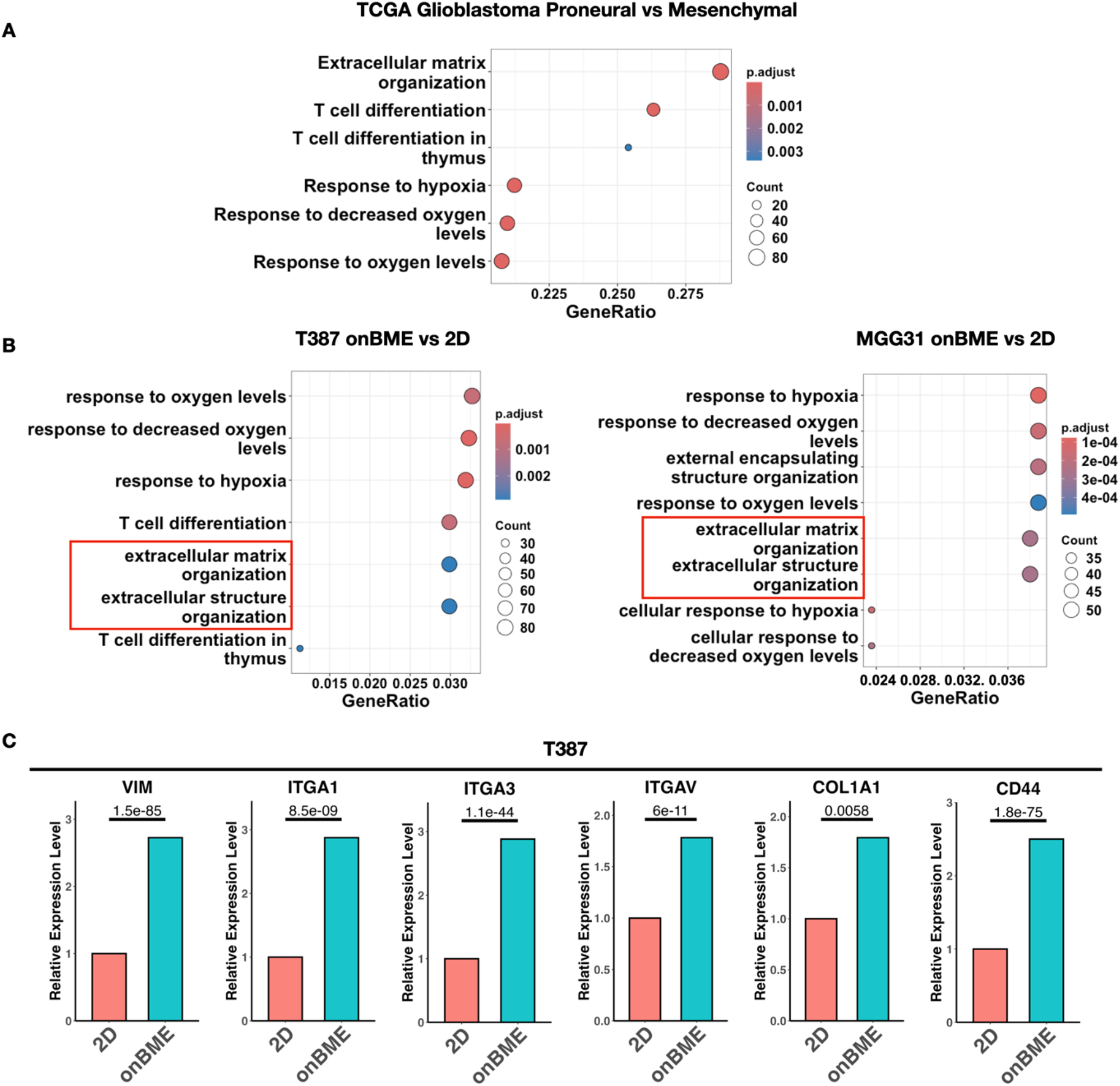
Culture onBME induces significant gene expression changes related to ECM and cell adhesion. **A**. GO enrichment of differentially expressed genes between the mesenchymal and proneural subtype of GBM in the TCGA dataset. **B**. GO enrichment of differentially expressed genes between the onBME and 2D culture of T387 and MGG31 mGBM cells. Extracellular matrix-related terms that are enriched for both cells are emphasized with red box. **C**. Relative expression level of important cytoskeleton, cell adhesion, and ECM component genes after culture onBME versus in 2D. p-values were calculated by DEseq2. N = 3.

We investigated if onBME culture induced more mesenchymal-like gene expression related to these processes. Performing similar enrichment on the DEGs comparing onBME versus 2D culture of mGBM cells showed that these same GO terms are among the most enriched (**Figure 2B-C and Supplementary Figure 2A**). This result indicates that onBME culture resulted in a more mesenchymal signature along the proneural-mesenchymal axis, including hypoxic metabolism and ECM remodeling.

We also tested specific genes related to known mesenchymal features (**Figure 2C and Supplementary Figure 2B**). Vimentin, the intermediate filament that is characteristic of mesenchymal cells, showed elevated expression levels of nearly threefold in T387 when cultured on BME. Certain integrin genes, such as ITGA1/3/V, have also been reported to be related to increased mesenchymal features, showing higher expression in mesenchymal versus proneural GBMs^15^. ITGA1, which binds to collagen type I as α_1_β_1_ has been shown to be upregulated during EMT^18,19^. Additionally, ITGA3 and V are both important markers for cancer cells undergoing EMT^20–22^. These integrin genes all showed increased expression in onBME cultures. ECM secretion is also an important feature for mesenchymal cancer cells and is believed to help facilitate their invasion^23^. Associated genes such as COL1A1, encoding collagen type I, and CD44, a surface glycoprotein that is an important mediator of cell-ECM interaction and a stemness marker of cancer cells, were also elevated^24^. The elevated expression of these important mesenchymal marker genes implied elevated mesenchymal feature of onBME cultures.

To discover potentially altered cellular processes of onBME cultures, we performed GSEA on the hallmarks gene sets. Interestingly, cell-cycle related gene sets, including G2M checkpoints, E2F targets, mitotic spindles, and MYC targets, all showed negative enrichment scores (**Supplementary Figure 3A-B**), consistent with smaller portion of proliferating cells in the mesenchymal subtype^15^. On the other hand, both cell lines were significantly enriched for hypoxia and differed in their metabolic pathways when cultured on BME, consistent with the GO enrichment analysis. T387 showed a negative enrichment score for oxidative phosphorylation (OXPHOS), implying decreased OXPHOS activity, which is in accordance with a positive hypoxia score. MGG31 showed a negative enrichment score for multiple lipid metabolic pathways, including cholesterol homeostasis, fatty acid metabolism, and adipogenesis (**Supplementary Figure 3A-B**). The distinct metabolic changes between T387 and MGG31 cultured on BME may have revealed different energy sources that can be used by mesenchymal GBM cells.

Collectively, these results revealed that the GBM onBME culture goes through important changes in gene expression towards signature mesenchymal cellular processes.

### OnBME culture induced mesenchymal-like cytoskeleton organization and morphology

Upregulation of genes involved in cytoskeletal organization and ECM remodeling in mGBM may promote the formation of cytoskeletal stress fibers, such as fibrous actin and intermediate filaments, rather than their localization at the cortical regions of the plasma membrane^25^. To determine whether onBME culture can represent this property, we examined the cytoskeletal organization of GBM cells under different culture conditions.

In onBME cell cultures, GBM cells exhibited well-organized actin stress fibers spanning the cell body along the axis direction, whereas cells cultured in 2D conditions displayed primarily cortical localization (**Figure 3A**). Vimentin, a key mGBM marker and intermediate filament protein, exhibits a pronounced polymerized fibrous structure in GBM cells onBME. In 2D culture, vimentin showed a reduced polymerized form, showing concentrated granules in T387 cells and bubble-like structures in MGG31 cells.

**Figure 3.**
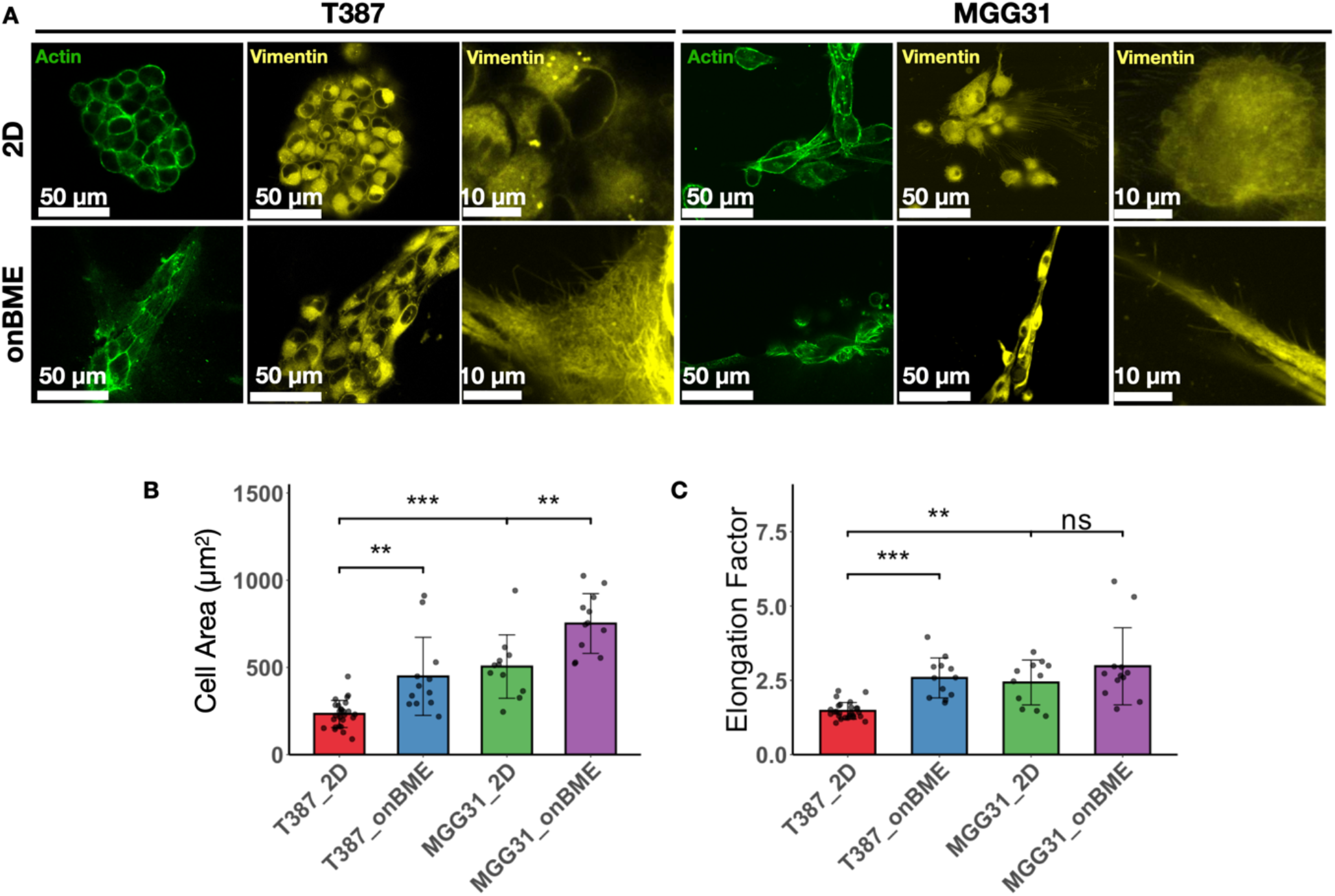
Culture onBME induces reorganization of cytoskeleton and cell morphology of mGBM cells. **A**. Actin and Vimentin in mGBM cells cultured in 2D or on BME2. **B**. Cell area quantification of mGBM cells under different culture condition. **C**. Elongation factor of mGBM cells under different culture condition. p-values were calculated using paired t-test. N = 11-20.

Different organization of the cytoskeleton regulates different cell morphology. We therefore quantified the cell area and elongation factor, defined as the ratio of the length of the longest to the shortest axis of a cell (**Figure 3B-C**). The results showed that both cell area and elongation are increased, reflecting increased mGBM features. Together, these results demonstrate that onBME culture promotes mesenchymal-like morphology and cytoskeleton organization, providing a more relevant model for studying mGBM cells.

### Co-culture of mGBM and microglia cells revealed enhanced recruitment of microglia when cultured on BME

Microglia cells are important immune cells frequently recruited to the GBM TME, where they contribute to tumor progression^26–28^. Notably, mGBM has been reported to exhibit increased recruitment of microglia cells^5,29^. We thus assessed whether the onBME model recapitulates the interaction between microglia and GBM cells, reshaping the TME.

Co-culture of microglia (HMC cells) and GBM cells onBME revealed that the two cell types formed an integrated network structure (**Figure 4A**). However, in 2D culture, T387 cells recruited microglia to the neurospheres, but the microglia cells remained individually to the surface of the spheres without observable integration and network formation. MGG31 cells showed simply mixture of two adherent cells without structural organization.

**Figure 4.**
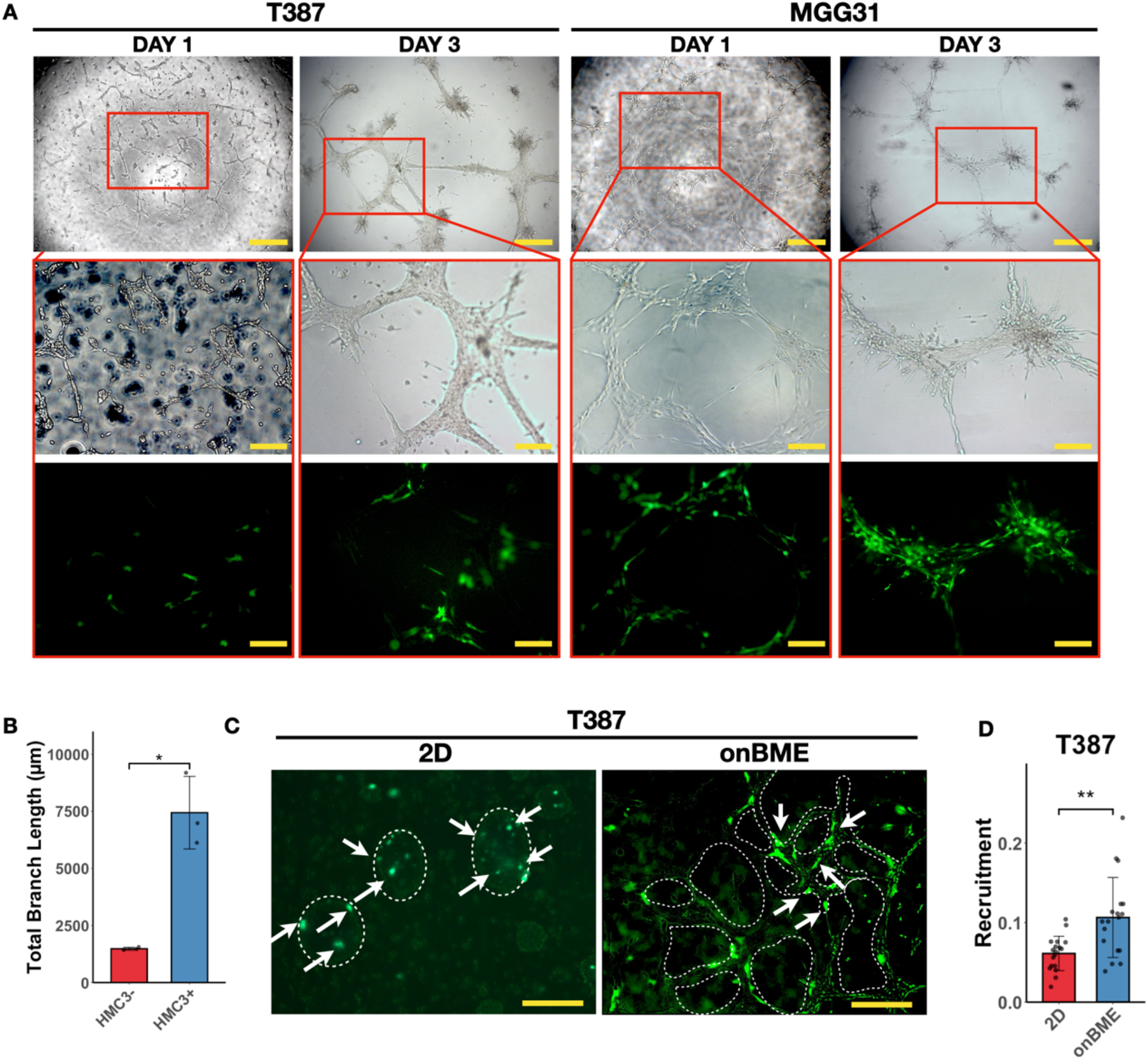
Co-culture of microglia and mGBM cells on BME shows increased recruitment. **A**. Integrated network structure formed by mGBM cells and microglia cell line HMC3 which expresses GFP. Scale bar = 500 µm for top panel and 200 µm for middle and bottom panels. **B**. Bar graph showing increased network formation quantified by total branch length after co-culture with microglia cells. P-value was calculated using t-test. N = 3. **C-D**. Comparing recruitment of microglia to the mGBM cell mass in 2D and on BME (C) and quantification (D). Representative tumor cell mass was outlined with dashed line and microglia cells expressing GFP are indicated with arrows. Scale bar = 200 µm. p-values were calculated using t-test. N = 18.

In addition, we characterized the network structure by quantifying total branch length and found that addition of microglia increased the connectedness of the GBM network over time (**Figure 4B**). Since network structure is thought to be a characteristic of invasiveness, the onBME model may reveal a potential contribution of microglia in promoting the motility of mGBM cells, although further experiments would be needed for confirmation.

Furthermore, we discovered that T387 cells on BME showed significantly increased recruitment compared to neurospheres in 2D (**Figure 4C-D**). These results indicate that onBME culture provides a more physiologically relevant model for the close interaction between mGBM and microglia cells, recapitulating increased microglia recruitment.

### OnBME model revealed repressed mGBM features by mGBM cells after CK2 inhibition

Due to the aggressive nature of mGBM, developing new therapeutic strategies to repress mesenchymal transition is crucial. Specifically, it is necessary to discover druggable targets that can suppress the transition. We hypothesized that the molecules highly activated onBME compared to 2D prompt GBM cells to adopt the mGBM subtype and serve druggable targets to suppress mesenchymal transition. Thus, we leveraged the Small Molecule Suit^11^ to identify potential drug targets that are highly expressed in GBM onBME compared to 2D. Among the top hits, CK2 emerged as a relatively underexplored target (**Supplementary Table 1**). Thus, we tested a panel of CK2 inhibitors, including CX4945, which is also known to bind a few more kinases^30^, and the two newly reported ones GO289^31^ and SGC-CK2-1^32^.

We performed RNA sequencing on CK2 inhibitor-treated cells to determine whether these compounds repress the molecular mGBM signature. GSEA analysis of DEGs comparing control and each drug-treated condition was conducted using two gene sets: the TCGA Mesenchymal versus Proneural gene set and the Hallmark EMT gene set. The results showed that treatment with CX-4945 and SGC-CK2-1 led to significantly negative enrichment scores, while GO289 also produced a negative—though less significant—enrichment score for the EMT gene set (**Figure 5A,B**). These findings suggest that GO289 can moderately, and CX-4945 and SGC-CK2-1 can significantly repress the mesenchymal signature in GBM cells.

**Figure 5.**
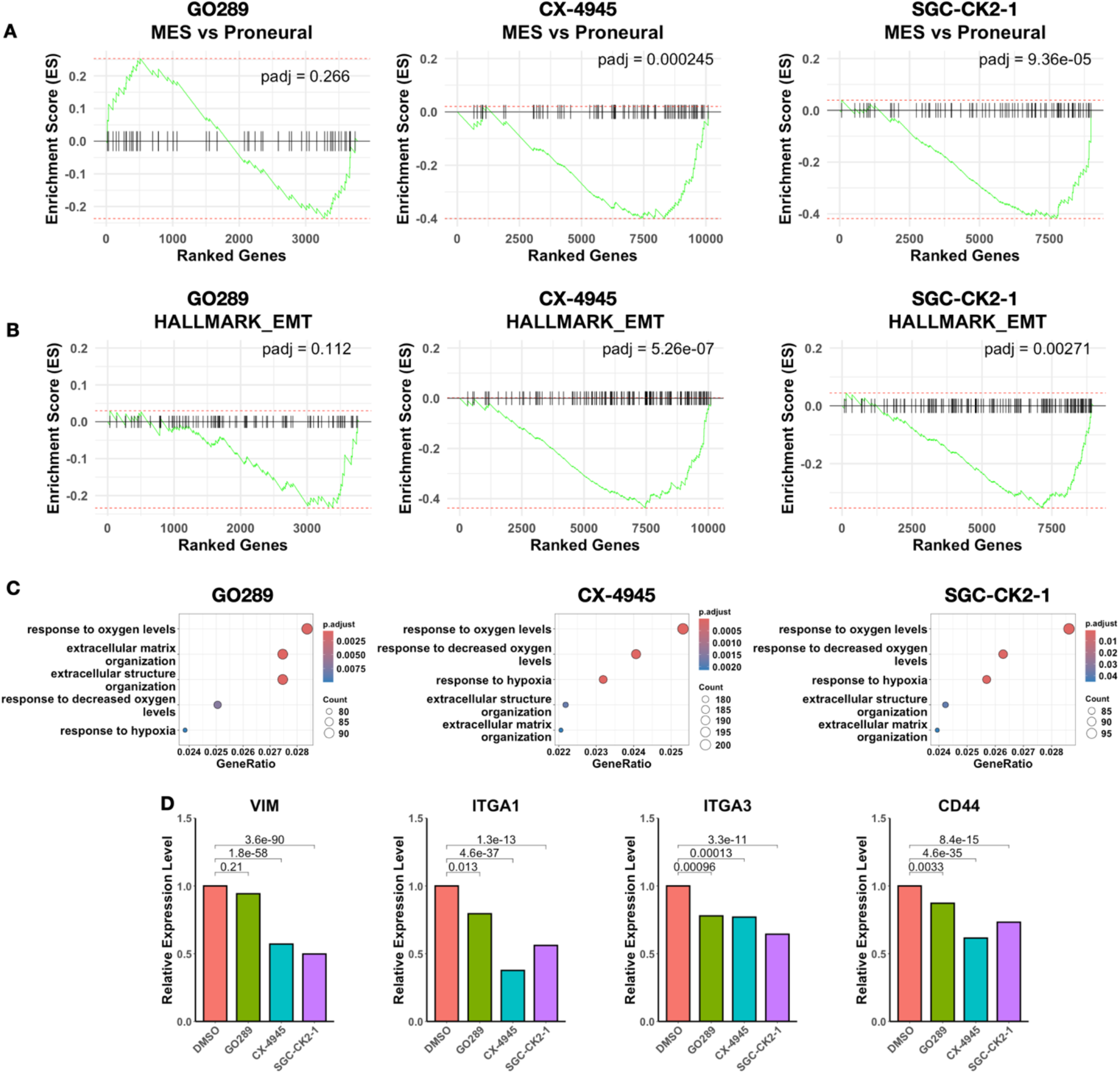
CK2 inhibitors repressed the mesenchymal signature of mGBM cells and related gene expression profiles. **A-B**. GSEA analysis of differentially expressed (DE) genes after treatment by CK2 inhibitors on the set of differential signature genes of mesenchymal versus proneural subtype (A) and on the hallmark epithelial to mesenchymal transition gene set (B). **C**. GO enrichment analysis of DE genes after CK2 inhibitors showed significant terms on extracellular matrix organization and response to hypoxia, both of which are associated with increased mesenchymal feature of mGBM cells as show in Figure 2. **D**. Bar graphs showing vimentin and mesenchymal-associated cell-ECM interaction genes are repressed by CK2 inhibitors. p-values were calculated by DEseq2. N = 3.

We also examined if mesenchymal-related cellular processes identified in the previous GO analysis are also repressed by the CK2 inhibitors. Indeed, all three drugs caused significant negative enrichment in the hypoxic metabolism genes and the ECM organization genes (**Figure 5C**). Similarly, key mGBM marker genes such as VIM, ITGA1/3, and CD44 are all downregulated following the CK2 inhibitors (**Figure 5D**). These results demonstrate that CK2 inhibitors can repress gene expression programs related to mGBM.

After confirming that CK2 inhibition reduced the gene expression signature of mGBM, we used the onBME model to test associated phenotypic changes. The CK2 inhibition reduced the network structures of GBM cells (**Figure 6A-B**). The extent of total branch length reduction after drug treatment is consistent with the extent of the repressed mesenchymal feature of each drug, with SGC-CK2-1 showing the most reduction and GO289 the least.

**Figure 6.**
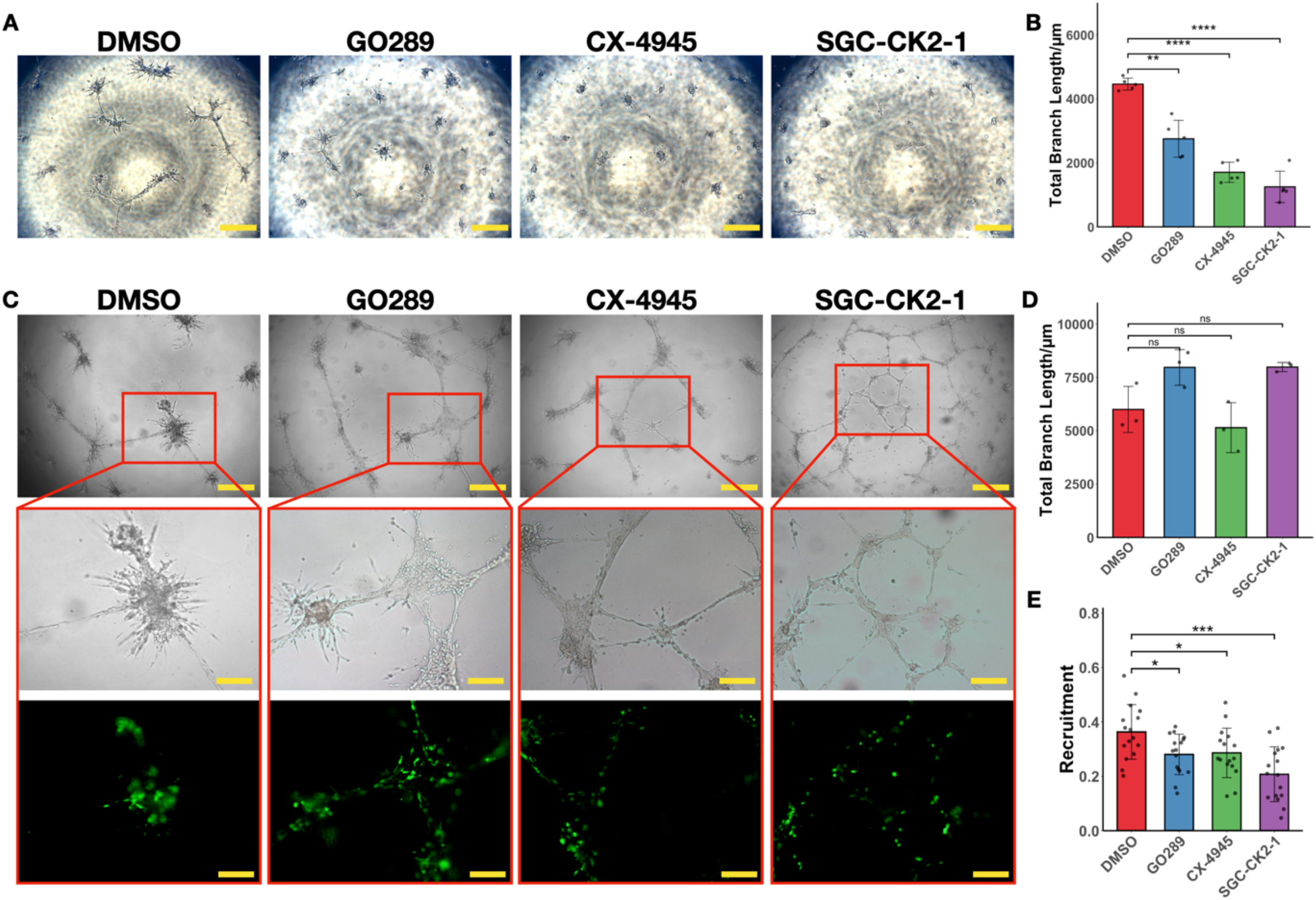
onBME model revealed repressed network-forming by CK2 inhibitors and reduced microglia recruitment. **A**. Bright filed images showing the morphology of networks formed by MGG31 cells onBME treated by either DMSO or CK2 inhibitors. Scale bar = 500 µm. **B**. Quantification of branch length of networks formed by MGG31 cells after CK2 inhibition. p-values were calculated by t-test. N = 5. **C**. CK2 inhibitor treatment on the co-culture of microglia with MGG31 on BME. Scale bar = 500 µm for top panel and 200 µm for middle and bottom panels. **D-E**. Bar graph showing maintained network connectedness after drug treatment (D), but recruitment of microglia cells is repressed by CK2 inhibitors(E). p-values were calculated by t-test. N=3 (D) or 26 (E).

We then investigated whether CK2 inhibitors affect the recruitment of microglia and their support of the network structure, reflective of mGBM invasiveness. The addition of microglia supported the network structure even under drug treatment, suggesting the role of microglia in promoting resistance to the drugs and sustaining mGBM invasiveness (**Figure 6C**). However, the recruitment of microglia was decreased, consistent with the ability of each drug to decrease mesenchymal features (**Figure 6D**). These findings further suggest that the CK2 inhibition attenuates the presentation of mGBM features, highlighting CK2 as a vulnerable node for targeting mesenchymal transition.

## Discussion

GBM remains one of the deadliest adult cancer types due to its aggressive nature and limited treatment options. The transcriptional subtypes of GBM cells underscore the importance of understanding subtype-specific biology when developing therapeutics. Among the different GBM subtypes, mesenchymal-like GBMs are associated with high stemness and the poorest prognosis. To better understand mGBM biology and enable therapeutic testing in a more physiologically relevant and accessible manner, we developed a gel-top model that enhances mGBM characteristics and recapitulates key features suitable for studying mGBM and evaluating drug susceptibility.

We found that GBM cells cultured on BME exhibit enhanced mGBM features compared to traditional 2D cultures. Gene expression associated with mesenchymal-related biological processes—such as reduced cell cycle progression and altered metabolism— was elevated. Additionally, onBME culture induced cytoskeletal and morphological changes consistent with a more mesenchymal-like phenotype. These findings suggest that the onBME system provides a more relevant model for studying the mesenchymal features of GBM cells.

Importantly, the onBME model is also compatible with the incorporation of other TME components, such as microglia, and is suitable for therapeutic testing. Using this model, we demonstrated enhanced recruitment of microglia toward GBM cells and a unique, integrated self-assembly pattern formed by mGBM and microglia cells, which represent mGBM features, not observed in 2D culture. These findings further support the relevance of onBME cuture for studying TME interactions in mGBM.

Furthermore, we identified CK2 as a potential drug target, representing genes that are highly expressed in onBME culture compared to 2D culture. We found that CK2 inhibitors suppress key mGBM features, including reduced expression of mesenchymal signature genes, inhibition of network formation by mGBM cells (a hallmark of invasiveness), and decreased recruitment of microglia. Interestingly, this model also revealed a previously unrecognized role of microglia in maintaining the network structure formed by mGBM cells.

In summary, culturing GBM cells on BME offers a novel and efficient platform for characterizing mesenchymal features. It is time-efficient and adaptable for various experimental and therapeutic testing purposes. This model provides a more physiologically relevant tool for future studies and drug screening in mGBM.

## Methods

### Cell culture

Patient derived glioblastoma cell line T387 was obtained from Dr. Jeremy Rich, and MGG31 was a gift from Dr. Hiroaki Wakimoto. GBM cells were cultured in complete neural media, which is composed of neurobasal medium (Gibco cat. 12348017), 1X B-27 supplement (Gibco cat. 17504044), 1X GlutaMax (Gibco cat. 35050061), 1X sodium pyruvate (Gibco cat. 11360070), 1% Penicillin-Streptomycin (Gibco cat. 15070063), 0.5 µg/mL FGF, and 0.5 µg/mL EGF. Microglia cell line HMC3 was purchased from ATCC (cat. CRL-3304) and maintained per ATCC protocol using EMEM media (ATCC 30-2003) containing 10% FBS. All cells were cultured at 37°C in 5% CO_2_.

For 2.5D culture, Cultrex Basement Membrane Extract Type 2 (BME2, biotechne cat. 3532-005-02), Corning Synthegel Spheroid matrix kit (Corning cat. 354789), and Vitrogel Hydrogel Matrix (The Well Bioscience cat. VHM01) were used per manufacturer’s protocol. In 96-well plates, 20 µL of matrix was coated on the bottom of each well, cured, and 10,000 resuspended single cells were added gently on top.

For 3D culture, cells were resuspended directly with pre-calculated amount of ice-cold BME2 to reach a density of 10,000 cell per 50 µL gel for one well of a 96-well plate. Gels containing cells were then added to the middle of each well to form a dom. Culture media were added to each well after the gels are solidified.

For co-culture of HMC3 and GBM cells, 2000 HMC3 cells were cultured with 10,000 GBM cells per well in 96-well plates in complete neural media.

### RNA extraction and sequencing

For RNA extraction experiments, cells were cultured in 24-well plates. For 2.5D onBME culture, 60 µL BME2 was used to coat each well and 200,000 cells were used. RNA extraction was performed one day after seeding the cells when full network structure was formed. Total RNA was extracted using TRIzol (Invitrogen cat. 15596026). Briefly, supernatant was carefully discarded, then 1mL of TRIzol was added to each well. The mixture was thoroughly homogenized by pipetting and transferred to a RNA-binding column (Zymo cat. C1004). The column was spin at 130,000 x *g* for 2 min, and the flow-through was discarded. Then the column was washed twice with washing buffer, and RNA was eluted in 30 µL nuclease-free water.

Library preparation from total RNA for RNA sequencing was done at *Azenta*. mRNA was selected from total RNA using poly-A selection. Paired-end sequencing was performed on an Illumina platform to deliver at least 30M reads per sample.

### Sequencing data analysis

Quality control of raw reads in *fastq* format was done using FastQC^33^, adaptor sequences was trimmed using trimgalore^34^. Reads that passed quality control were mapped to human genome hg38 using hisat2 paired-end mode with default parameters. Feature counting was done using featureCount function from the subread^35^ package. Differentially expressed genes were found using DEseq2^36^. GSEA of differentially expressed genes ranked by log2 fold change was done using the fgsea^37^ package in R with default parameters. Enrichment analyses for gene ontology terms were done using the clusterProfiler^38^ package in R.

The MES vs Proneural gene set was defined by finding differentially expressed genes comparing the mesenchymal and proneural subtype of TCGA dataset using R2 suit. To define a gene set with optimal size for GSEA, we set a stringent threshold of log2 fold change < -2 and p-value less than 0.001. to filter the differentially expressed genes.

### Cytoskeleton staining and imaging

For fluorescent imaging, cells were cultured in ibidi µ-Slides (ibidi cat. 81507). For onBME culture, 10µL BME2 was used for each well. Actin and vimentin were stained using SPY555 (Cytoskeleton cat. CY-SC202) and BioTracker TiY Vimentin live cell dye (Sigma cat. SCT059). Manufacturer-recommended concentration was used. Dyes were added to culture media and incubated for an hour before imaging in live cell. Imaging was done on a Leica MICA platform. Cell area and elongation factor was quantified using a customized Matlab script.

### Generation of GFP-expressing HMC3 cells

GFP-expressing HMC3 cells were generated by lentiviral transduction of a pCDH-GFP vector. Lentivirus was produced by lipofectamine 3000 (Invitrogen cat. L3000015) transfection of the envelop, packaging, and expression vector into HEK-293T cells per manufacturer’s protocol. Transduction was done when HMC3 cells reached 30% confluency, 2% virus and 5 µg/mL polybrene (Sigma cat. TR-1003-G) was added to the cells, and the plate was centrifuged at 2000 rpm for 20 minutes. After 48 hours of transduction, 10 µg/mL puromycin was added and selection was done for five days, with culture media changed every two days.

### Quantification of microglia recruitment

We quantified the ratio of the GFP-expressing area, representing recruited GFP-labeled HMC3 microglia cells, within the neurosphere or network relative to the total area of the neurosphere or network, using a custom-made Matlab (Mathwork, MA) algorithm.

### Small molecule suite analysis

To find potential targets that can repress the mesenchymal feature of GBM cells, we queried the up-regulated genes when MGG31 is cultured on BME in the small molecule suite^11^. We retained drugs that are chose for their selectivity and in early clinical trial stage (I and II) and found that CK2 is among the most sensitive targets.

### Drug treatment

GO289 was a gift from the lab of Dr. Tsuyoshi Hirota. CX-4945 (MCE cat. HY-50855) and SGC-CK2-1 (MCE cat. HY-139004) was purchased from MedChemExpress. Media containing working concentration of drugs (2.5 µM for GO289 and CX-4945, 1.25 µM for SGC-CK2-1) was used to resuspend cells for seeding. Images of drug-treated culture was taken 72 hours after seeding.

## Acknowledgements

This work was supported by the following funds: CHLA start-up support (YP, and JP), NIH grant CA238662 (JNR and SAK).

## Author contributions

Yuanzhong Pan: Data curation, data analysis, writing of manuscript, review and editing

Priscilla Chan: Data curation

Jeremy Rich: Conceptualization, supervision, funding acquisition, review

Steve Kay: Conceptualization, supervision, funding acquisition, review, and editing

JinSeok Park: Conceptualization, supervision, funding acquisition, data analysis, writing, review, and editing.

## Data and Code Availability

Raw sequencing data in fastq format and processed counts data are available on GEO through accession number GSE292931. Codes used for analyzing the data in R can be found in the Github repository https://github.com/yzpan1/glioblastoma-onBME.

## Figure legends

**Supplementary Table 1**.

List of up-regulated genes when MGG31 are culture on BME compared to 2D and small molecule suite query results.

---

**Supplementary Figure 1.**
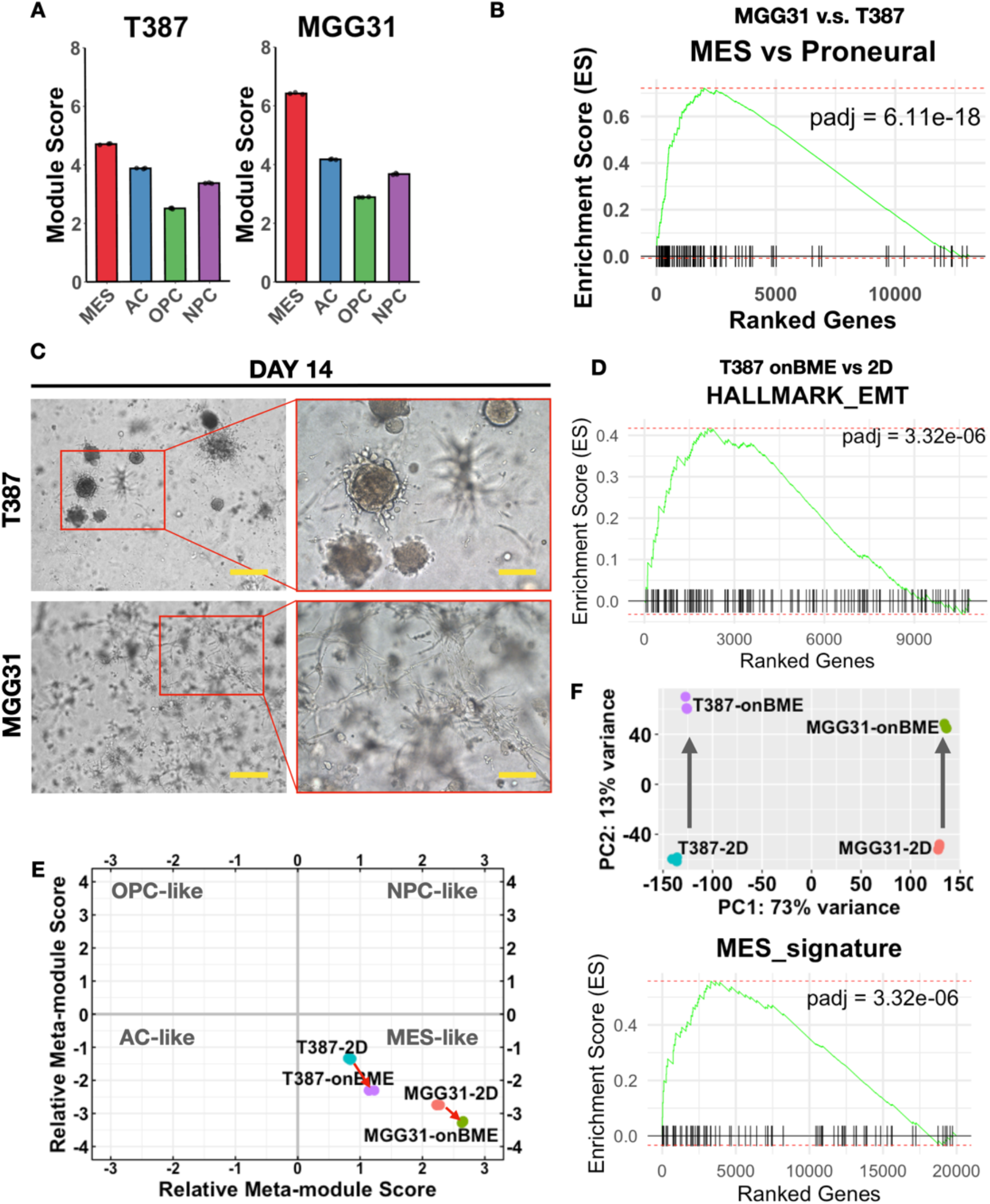
**A**. Bar graph showing module scores of two GBM cell lines, confirming their mesenchymal subtpe. **B**. GSEA analysis of the differentially expressed genes between MGG31 and T387 on the mesenchymal versus proneural gene set showed significant enrichment. **C**. Images of the mGBM cells cultured in 3D (inside BME2) after 14 days of seeding. Scale bar = 500 µm in original image and 200 µm in enlarged image. **D**. GSEA analysis of differentially expressed genes comparing 2D and onBME culture of T387 cells shows increased EMT feature. **E**. Two-dimensional representation of GBM subtypes showing that both mGBM cell lines exhibit higher mesenchymal signature after onBME culture. **F**. PCA analysis showing that PC2 accounts for most of the variance between 2D and onBME culture (top), and GSEA analysis on the components of PC2 showing significant enrich score for the MES signature gene set.

**Supplementary Figure 2.**
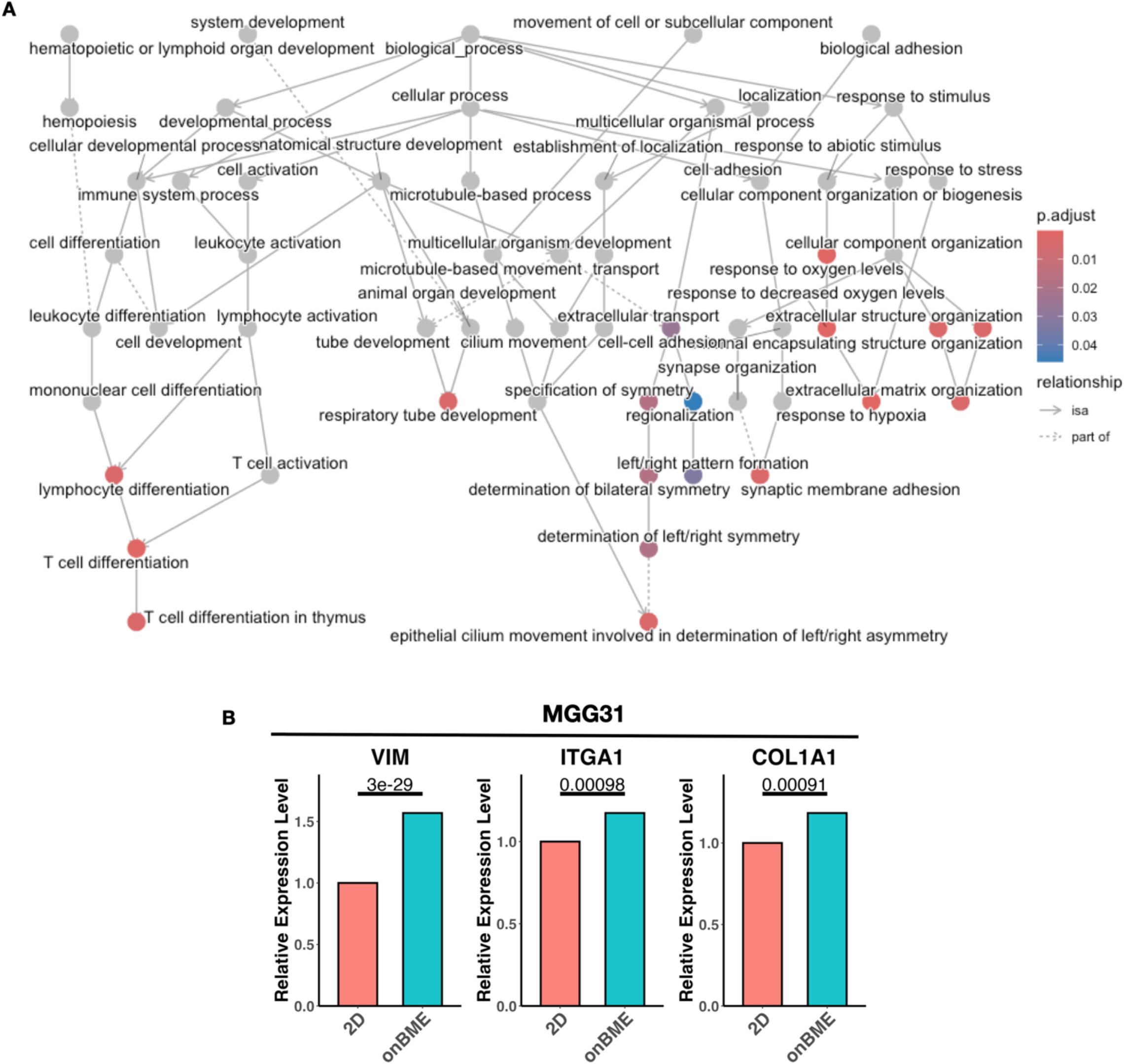
**A**. Full GO enrichment plot of differentially expressed genes of T387 cell cultured on BME2 versus 2D. **B**. Bar graph showing increased vimentin, integrin and collagen genes in MGG31 comparing onBME to 2D culture. p-values were calculated by DEseq2. N = 3.

**Supplementary Figure 3.**
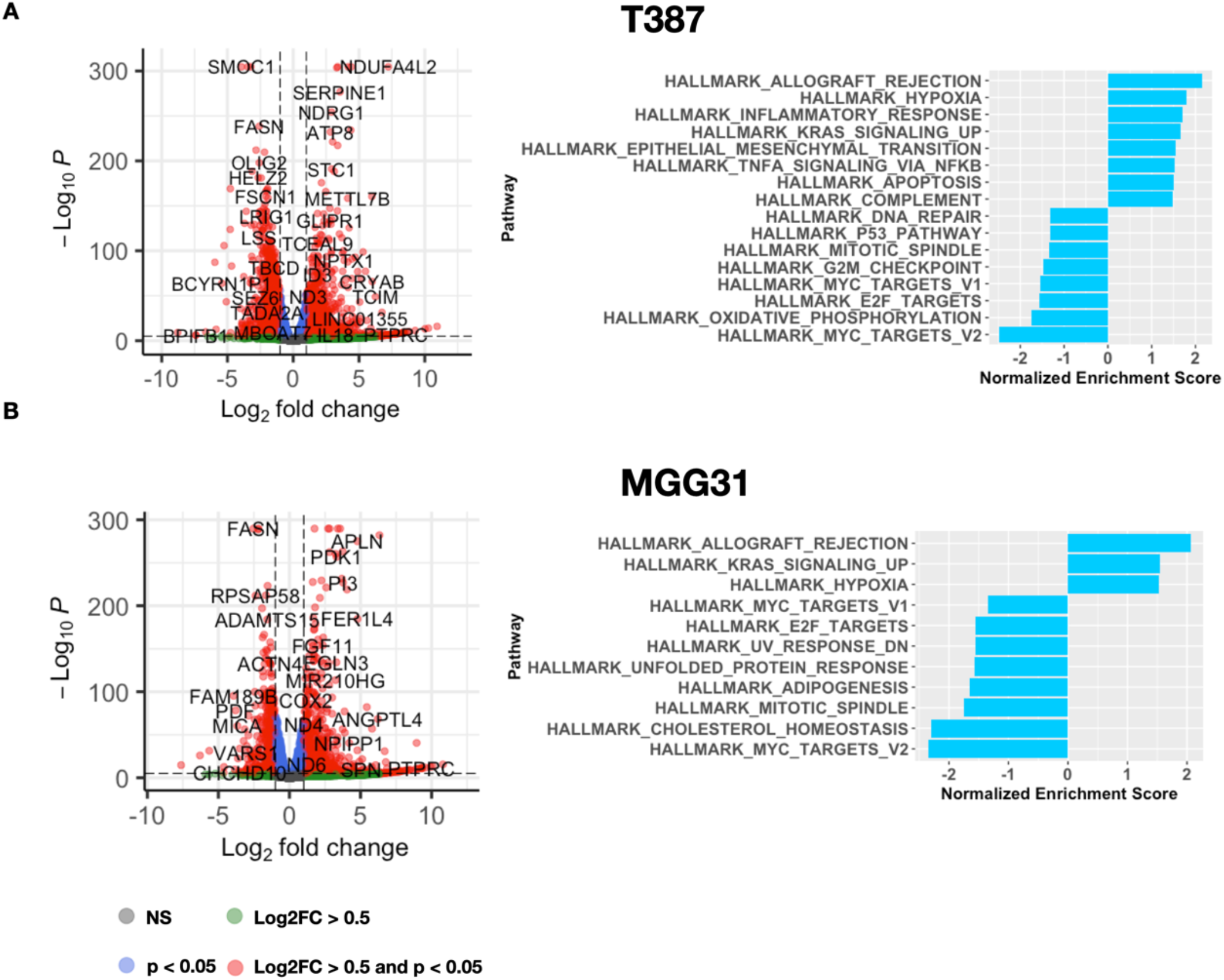
**A-B**. Volcano plot of differentially expressed genes comparing onBME and 2D culture of T387 (A) and MGG31 (B) cells, and GSEA analysis of DE genes on hallmark pathways. Only significantly enriched pathways are plotted.

